# Certified Design Regions under Parametric Plant Uncertainty with Application to Antithetic Integral Feedback

**DOI:** 10.64898/2026.10.08.757569

**Authors:** Rudra Prakash, Shaunak Sen

## Abstract

Kinetic parameters in engineered biomolecular circuits are typically known only within intervals and are not precisely tunable; controller synthesis is therefore naturally posed over sets of gains rather than at a single nominal design. We formulate robust controller tuning under parametric plant uncertainty as a universally quantified set-inversion problem: given a plant-parameter box, compute the controller gains that stabilize every plant in the box. Robust stability is decided by Hurwitz sign conditions, which are bounded over the joint gain–parameter domain using validated range enclosures, yielding machine-checkable certificates of local exponential stability of the deterministic equilibrium. Numerical experiments show that the certified region depends strongly on the range enclosure. For an antithetic integral feedback (AIF) controller acting on a two-species gene-expression cascade, a tensor-product Bernstein enclosure certifies 83.2% of the design plane at bisection tolerance *ε* = 0.05 with 324 terminal boxes, whereas four baseline enclosures certify between 50.9% and 58.9% on the same paving. For a fifth-order loop, the certified fraction increases from 34.4% to 73.8%, and the certified and excluded terminal boxes confine the robustly stabilizing area to a window of about 2.1 percentage points. Finally, for a refining variant of the algorithm in which the plant box is refined alongside the design box, the undecided area satisfies |*U* (*ε*)| = *O*(*ε*) under a non-degeneracy assumption on the stability boundary.

## 1 Introduction

Controllers built from biomolecular parts must operate with kinetic parameters that are uncertain and context dependent. Degradation, translation, and sequestration rates are often reported only to within multiplicative factors and can drift with growth conditions and host context. As a consequence, controller gains that stabilize a nominal model may fail to stabilize plants within a prescribed uncertainty set. The antithetic integral feedback (AIF) motif, which provides robust perfect adaptation in synthetic biology [1, 2], serves as a representative example: while its nominal stability and performance limitations have been characterized in detail [3], gain choices that stabilize a nominal plant need not be robust to parametric uncertainty.

A standard response is to perform sensitivity sweeps or Monte Carlo simulations. Such studies provide evidence but not guarantees, since absence of observed instability on a finite sample does not imply stability over a continuous uncertainty set. The objective of this paper is to provide certified alternatives to sampling-based acceptance rules: over a continuous gain domain, we compute guaranteed inner and outer approximations of the gain set that stabilizes every plant in a specified parameter box.

Interval analysis provides a natural framework for such certification [4, 5]. Related validated-numerics tools have recently been applied to uncertain biomolecular circuits for equilibrium enclosure [6] and certification of bifurcation candidates [7], with complexity analyses of the underlying solvers in [8]. Those results address *analysis* problems, in which the circuit is fixed and one certifies properties of the dynamics over an uncertainty box. The present paper addresses a *synthesis* problem: given an uncertainty box and a specification, compute and certify the set of controller parameters satisfying the specification, together with a quantitative account of the remaining undecided region at a prescribed bisection tolerance.

Beyond validated numerics, there is a line of work on characterizing controller parameter regions that meet stability or performance specifications. Ackermann’s parameter-space approach [9] maps eigenvalue specifications into the controller-parameter plane and traces stability boundaries. In parametric robust control, Kharitonov’s theorem [10] provides effective sufficient conditions for robust stability of interval polynomial families; it applies whenever the characteristic coefficients can be enclosed in intervals, but it treats them as independent, whereas in biomolecular loops they usually depend on the same few parameters; we use it as a baseline to measure what this costs.

Bernstein expansion is another classical tool for robustness *analysis* under polynomial parameter dependence [11, 12], typically used to decide stability of a prescribed uncertain family. Interval methods have also been applied to controller tuning by bounding robustness and performance measures over uncertainty sets [13]. In contrast, the present work returns an explicit set approximation over a continuous design domain, partitioned into certified-stabilizing, certified-excluded, and undecided boxes, which yields two-sided area brackets and supports direct comparison with sampling-based acceptance rules. Sums-of-squares programming [14] can likewise certify positivity over the joint gain–plant-parameter domain, but it typically requires proposing a candidate region and increasing the relaxation degree until feasibility is achieved; set inversion instead produces an explicit approximation of the admissible set together with an explicit undecided region. From the viewpoint of real algebraic geometry, the robust design set is semi-algebraic, defined by polynomial inequalities with a universal quantifier. Here we seek certified numerical approximations rather than a symbolic characterization.

Set inversion has been used to compute stability domains since Walter and Jaulin [15], who obtained guaranteed inner and outer approximations of the stable region in parameter space. The quantifier structure here is different: [15] characterizes those parameters for which a fixed polynomial is stable, whereas the robust design set *K* requires stability for every plant in an uncertainty box. This universal quantifier over **p** motivates the asymmetric exclusion rule in Proposition 1. The closest prior work is [16], which uses interval bisection and constraint propagation to design parameter intervals meeting steady-state specifications in biomolecular circuits and identifies stability certification as future work; the present paper addresses that step.

The main contributions of the paper are as follows:

1. We formulate controller tuning under parametric uncertainty as a universally quantified set-inversion problem, *K* = *{****κ*** : ∀ **p** ∈ [**p**], the closed-loop Jacobian is Hurwitz*}*, with robust stability expressed through finitely many polynomial sign conditions. We develop sound inner and outer classification tests, yielding a subpaving of the design domain into certified-stabilizing, certified-excluded, and undecided boxes.
2. We construct tensor-product Bernstein inclusion functions for these sign conditions and incorporate them into the quantified paving. Bernstein enclosures are classical [11, 12]; the focus here is on the effect of the enclosure choice on the certified design region, assessed against four baselines (natural interval extension, centered form, natural extension of the unexpanded Routh expression, and a Kharitonov test on the coefficient box) on an identical paving, design domain, witness set, and stopping rule.
3. We establish an asymptotic completeness statement for a refining variant of the algorithm: when the plant box is refined alongside the design box, the undecided area satisfies |*U* (*ε*)| = *O*(*ε*) under a non-degeneracy assumption on the stability boundary (Proposition 3).
4. We report numerical results on two AIF loops of fourth and fifth order. In both loops, part of the robust stability boundary at large sequestration rates admits a closed-form expression, which is used to corroborate the certified region. Representative terminal boxes are also verified in exact rational arithmetic.

### Paper organization

Section 2 states the robust design problem and introduces the AIF loop used as a running example. Section 3 develops the certified inclusion tests and the Bernstein enclosure, and Section 4 summarizes the resulting paving algorithm. Section 5 reports numerical experiments on the two loops, and Section 6 concludes.

#### Notation

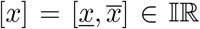 denotes a real interval and [**x**] an interval vector (box), of width 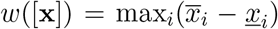. For *f* : ℝ^*n*^ → ℝ, an *inclusion function* [*f*] satisfies *f* ([**x**]) ⊆ [*f*]([**x**]) for every box [**x**]. | *·* | denotes Lebesgue measure and *D* the design domain used in the numerical experiments.

## 2 Problem Formulation

### 2.1 The robust design set

Let 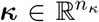 denote the design parameters of a closed loop and let **p** denote the uncertain plant parameters, assumed to lie in a box [**p**]. We assume that the closed loop admits a known positive equilibrium and that, on the domain of interest, the characteristic polynomial of the Jacobian at that equilibrium has positive coefficients. Under this assumption, the Hurwitz (equivalently, Liénard– Chipart) criterion reduces local exponential stability to finitely many sign conditions *g*_*j*_ (***κ*, p**) *>* 0, *j* = 1, …, *m*, where each *g*_*j*_ is a Hurwitz determinant evaluated at the equilibrium. In the examples considered below, each *g*_*j*_ is a rational function of the parameters.

#### Definition 1

(Robust design set). 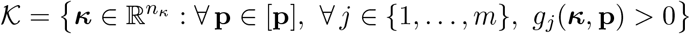.

**Remark 1**. *The set K is not a neighborhood of a nominal stabilizing gain. Rather, it collects those gains that satisfy the stability inequalities uniformly over the full uncertainty box* [**p**]; *consequently, gains that stabilize a nominal plant may lie outside K. Section 5 compares nominal-parameter tuning with robust stability on the design domains considered*.

We seek to compute inner and outer approximations 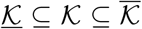 consisting of boxes, together with certificates.

### 2.2 Running example: antithetic integral feedback

We consider the AIF controller of [1] acting on a two-species gene expression cascade. With controller species *z*_1_, *z*_2_ and plant species *x*_1_, *x*_2_, the closed-loop dynamics are

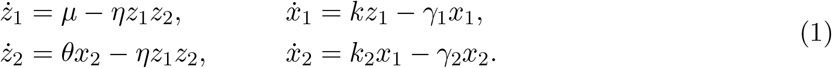

Subtracting the two controller equations yields *ż*_1_ −*ż*_2_ = *µ* −*θx*_2_, which provides integral action. At any positive equilibrium, this implies 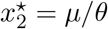, independently of the plant parameters. For positive parameters, the remaining equilibrium components are uniquely determined, and (1) admits a unique positive equilibrium,

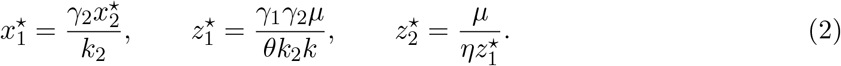

Define 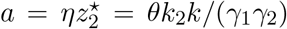 and 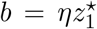, so that *ab* = *ηµ*. The Jacobian at (2), in the ordering (*z*_1_, *z*_2_, *x*_1_, *x*_2_), is

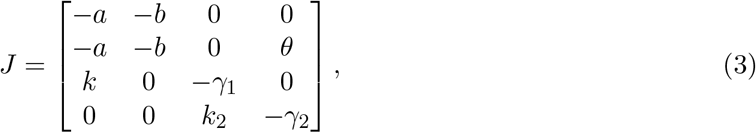

whose characteristic polynomial *s*^4^ + *a*_1_*s*^3^ + *a*_2_*s*^2^ + *a*_3_*s* + *a*_4_ has

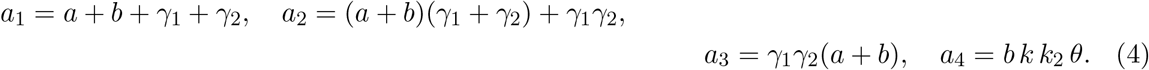

#### Lemma 1.

*For all positive parameter values, a*_1_, *a*_2_, *a*_3_, *a*_4_ *>* 0. *Consequently, J in* (3) *is Hurwitz (and* (1) *is locally exponentially stable at* (2)*) if and only if the third Hurwitz determinant*

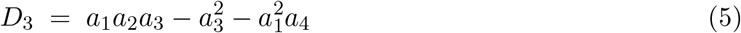

*is positive*.

*Proof*. Each coefficient in (4) is a sum of monomials with positive factors, hence *a*_*i*_ *>* 0 for *i* = 1, …, 4. For a monic quartic, the Hurwitz criterion is equivalent to the inequalities *a*_1_ *>* 0, *a*_1_*a*_2_ − *a*_3_ *>* 0, *D*_3_ *>* 0, and *a*_4_ *>* 0. It remains to verify that *a*_1_*a*_2_ − *a*_3_ *>* 0 holds on the positive orthant. Writing *σ* = *a* + *b* and Γ = *γ*_1_ + *γ*_2_, one obtains *a*_1_*a*_2_ − *a*_3_ = *σ*^2^Γ + *σ*Γ^2^ + *γ*_1_*γ*_2_Γ, which is strictly positive for *σ*, Γ, *γ*_1_, *γ*_2_ *>* 0. Therefore, among the Hurwitz inequalities, only *D*_3_ *>* 0 can be violated.□

Lemma 1 reduces the stability test to a single scalar inequality. We take ***κ*** = (*η, k*) as the design parameters, corresponding to the sequestration rate and the actuation gain, and treat them as tunable quantities. The uncertain plant parameters are **p** = (*γ*_1_, *γ*_2_, *k*_2_). Writing *D*_3_(***κ*, p**) for (5) evaluated using (4) and (2), Definition 1 applies with *m* = 1 and *g*_1_ = *D*_3_.

The robust design problem is nontrivial at fourth order. With a single plant species 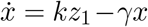 and sensing of *x*, the closed loop is third order; in that case the only nontrivial Hurwitz inequality expands to *a*_1_*a*_2_ − *a*_3_ = *γ*(*a*^2^ + *ab* + *b*^2^) + *γ*^2^(*a* + *b*) *>* 0, so every positive gain pair is stabilizing.

In addition, part of the stability boundary of (1) can be derived in closed form. Setting *µ* = *θ* = 1, writing *P* = *γ*_1_*γ*_2_ and using *σ* = *a* + *b* and Γ = *γ*_1_ + *γ*_2_ as in the proof of Lemma 1, one obtains *a*_3_ = *Pσ, a*_4_ = *ηP*, and

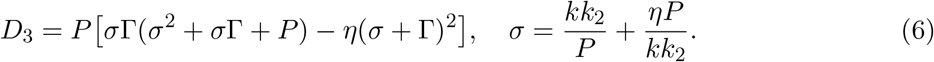

For fixed *k* and *η* → *∞*, both terms in the bracket scale as *η*^3^. It follows that *D*_3_ *>* 0 for all sufficiently large *η* if *kk*_2_ *< γ*_1_*γ*_2_(*γ*_1_ + *γ*_2_), whereas *D*_3_ *<* 0 for all sufficiently large *η* when the inequality is reversed. For the plant box of Section 5, this yields the robust threshold log_10_ *k <* −0.457, attained at the corner (*γ*_1_, *γ*_2_, *k*_2_) = (0.8, 0.4, 1.1); the corresponding nominal threshold is log_10_ *k <* −0.125. Section 5 uses these values to corroborate the certified region independently of interval arithmetic.

This asymptotic argument identifies only one portion of the boundary (the upper edge of the unstable set at large *η*). It does not characterize the dependence of that edge on *η*, nor the value of *η* below which the unstable set disappears. The certified map of Section 5 gives the remainder of the boundary, together with a guaranteed range for the size of the robust region.

## 3 Certified Test

### 3.1 Inner and outer tests

Set inversion [4] bisects the design plane and classifies each box [***κ***] by an inclusion test; the quantifier over **p** is absorbed as follows.

#### Proposition 1

(Soundness). *Let* [*g*_*j*_] *be an inclusion function for g*_*j*_ *on* [***κ***] *×* [**p**], *j* = 1, …, *m*.

*If* [*g*_*j*_]([***κ***], [**p**]) *>* 0 *for every j, then* [***κ***] ⊆ *K*.

*If there exist* **p**^*†*^ ∈ [**p**] *and an index j such that* 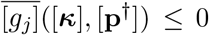, *where* [**p**^*†*^] *denotes the degenerate box at* **p**^*†*^, *then* [***κ***] ∩ *K* = ∅.

*In the running example, m* = 1 *and one may take g*_1_ = *D*_3_ *or, equivalently on the considered domain, g*_1_ = *q (Section 3*.*2)*.

*Proof*. (1) By the inclusion property, *g*_*j*_ ([***κ***] *×* [**p**]) ⊆ [*g*_*j*_]([***κ***], [**p**]) for each *j*. If [*g*_*j*_]([***κ***], [**p**]) *>* 0, then *g*_*j*_ (***κ*, p**) *>* 0 for all (***κ*, p**) ∈ [***κ***] *×* [**p**]. If this holds for every *j*, then [***κ***] ⊆ *K* by Definition 1.

(2) Let **p**^*†*^ ∈ [**p**] and let [**p**^*†*^] denote the corresponding degenerate box. If 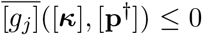 for some index *j*, then *g*_*j*_ (***κ*, p**^*†*^) ≤ 0 for all ***κ*** ∈ [***κ***]. Consequently, no ***κ*** ∈ [***κ***] can satisfy *g*_*j*_ (***κ*, p**) *>* 0 for all **p** ∈ [**p**], and hence [***κ***] ∩ *K* = ∅.

The two conditions of Proposition 1 are intentionally asymmetric. In particular, exclusion does not require a non-positive enclosure over the full plant box, which is typically conservative; it suffices to exhibit a single witness plant **p**^*†*^ ∈ [**p**] for which at least one stability inequality fails for every gain in the design box. More generally, the witness set may be prescribed arbitrarily and the exclusion test does not rely on monotonicity of *g*_*j*_ with respect to **p**.

### 3.2 Bernstein range bounds

The effectiveness of the inner test depends on the tightness of the range enclosure used to decide the sign of *g*_*j*_ on [***κ***] *×*[**p**]. For the running example, substituting (2) into (4) and clearing denominators yields 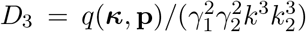. Since the denominator is a strictly positive monomial on the domain of interest, *D*_3_(***κ*, p**) *>* 0 if and only if *q*(***κ*, p**) *>* 0, and we apply Proposition 1 to the polynomial *q*.

The polynomial *q* has 24 terms, with multidegree (7, 7, 6) in (*γ*_1_, *γ*_2_, *k*_2_) and (3, 6) in (*η, k*). The natural interval extension of *q* evaluates repeated variable occurrences independently and can therefore be conservative on [**p**], since *γ*_1_, *γ*_2_, and *k*_2_ appear at high degree and in many terms.

We instead bound *q* using its Bernstein coefficients. For a box 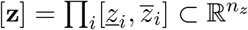, represent *q* in the tensor-product Bernstein basis of multidegree *d*. The Bernstein coefficients *{b*_*α*_*}* satisfy the range-enclosure property

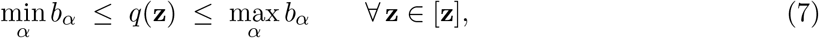

and the resulting enclosure sharpens quadratically under subdivision [12].

#### Proposition 2.

*Define* [*q*]_*B*_([**z**]) := [min_*α*_ *b*_*α*_, max_*α*_ *b*_*α*_]. *Then* [*q*]_*B*_ *is an inclusion function for q, and combining it with Proposition 1yields a sound inner test*.

*Proof*. By (7), *q*([**z**]) ⊆ [*q*]_*B*_([**z**]), so [*q*]_*B*_ is an inclusion function. Soundness of the corresponding inner test follows from Proposition 1(1).□

The construction in Proposition 2 does not depend on the particular form of *q*. It applies to any rational condition *g*_*j*_ whose denominator is positive on the domain and can be cleared. With multiple conditions, the inner test requires all Bernstein lower bounds to be positive. For simplicity, Proposition 3 below is stated for a single condition, as in the running example.

The affine map to [**p**] and the Bernstein transform act one axis at a time. Accordingly, the transform along the three plant axes can be computed once over [**p**] and reused for every design box; only the two design axes are transformed per box.

### 3.3 Completeness

Soundness alone does not imply that the undecided region vanishes under refinement. Define *φ*(***κ***) = min_**p**∈[**p**]_ *q*(***κ*, p**) so that *K* = *{****κ*** : *φ*(***κ***) *>* 0*}*. Let *B*(***κ***) denote the Bernstein lower bound for *q*(***κ***, *·*) computed over the full plant box [**p**] at a fixed design point ***κ***; by construction *B*(***κ***) ≤ *φ*(***κ***). The following proposition gives a sufficient condition under which a refining variant of the algorithm yields an undecided area that decays linearly with the bisection tolerance *ε*.

#### Proposition 3

(Completeness with a linear rate). *Run Algorithm 1in its* refining *form: certify* [***κ***] *when* [*q*]_*B*_ *>* 0 *on* [***κ***] *×* [**p**]_*i*_ *for every member of a dyadic cover {*[**p**]_*i*_*} of* [**p**] *of width at most ε, and exclude it when* 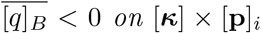 *for some member. Assume the level set {φ* = 0*} is non-degenerate, in the sense that* |*{****κ*** ∈ *D* : |*φ*(***κ***)| ≤ *t}*| = *O*(*t*) *as t* → 0. *Then*

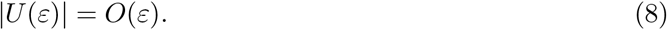

*Proof*. All computations are carried out in the bisected coordinates (log_10_ *η*, log_10_ *k*). The change of variables to (*η, k*) is bi-Lipschitz on the compact design domain, so bounds expressed in terms of the box width are preserved up to constant factors.

Since *q* is a polynomial, it is Lipschitz on any compact set; let Λ be a Lipschitz constant of *q* on the relevant subset of (***κ*, p**)-space. Moreover, by the standard Bernstein range-overestimation estimate under subdivision [12], there exists a constant *C >* 0 (depending only on the compact domain and the chosen multidegree) such that for any box *X* of width at most *ε*, [*q*]_*B*_(*X*) ≥min_*X*_ *q* − *Cε*^2^ and 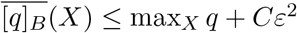.

Let [***κ***] be undecided and let ***κ***_*c*_ denote its center. Since [***κ***] is not certified, there exists an index *i* such that [*q*]_*B*_ ≤ 0 on *X* = [***κ***] *×* [**p**]_*i*_, and therefore min_*X*_ *q* ≤ *Cε*^2^. Choose (***κ***^*′*^, **p**^*′*^) ∈ *X* attaining the minimum. Then *φ*(***κ***_*c*_) = min_**p**∈[**p**]_ *q*(***κ***_*c*_, **p**) ≤ *q*(***κ***_*c*_, **p**^*′*^) ≤ *q*(***κ***^*′*^, **p**^*′*^) + Λ*ε* ≤ *Cε*^2^ + Λ*ε*. Since [***κ***] is not excluded, let **p**^*⋆*^ be a minimizer of *q*(***κ***_*c*_, *·*) over [**p**] and let 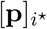 be a member of the dyadic cover containing **p**^*⋆*^. If *φ*(***κ***_*c*_) *<* −(2Λ*ε* + *Cε*^2^), then for all 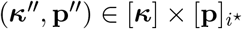,

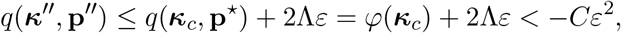

which implies 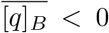 on 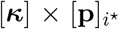 and would exclude [***κ***], a contradiction. Hence *φ*(***κ***_*c*_) ≥−(2Λ*ε* + *Cε*^2^).

Combining the two inequalities yields |*φ*(***κ***_*c*_)| ≤ *Kε* with *K* = 2Λ + *Cε*. Thus, the center of every undecided box lies in a *O*(*ε*)-neighborhood of the level set *{φ* = 0*}*. Since *φ* is Lipschitz (as the pointwise minimum of the Lipschitz family *q*(***κ***, *·*)), each undecided box of width at most *ε* is contained in a band of the form *{*|*φ*| ≤ *cε}* for a suitable constant *c*. The non-degeneracy hypothesis then implies |*U* (*ε*)| = *O*(*ε*).□

The hypothesis concerns the geometry of the level set *{φ* = 0*}* rather than the tightness of the Bernstein bound over the unrefined plant box; in particular, no assumption is made on the set *{B >* 0*}*. The statement is formulated in exact arithmetic. In finite precision, the enclosure error saturates at the rounding level, so (8) describes the regime above that saturation. The numerical certificates reported in Section 5 are unaffected, since each is verified at the tolerance at which it is computed.

#### Remark 2

(Cost of the refinement). *Without refinement of the plant box, the certified region converges to {B >* 0*}* ⊆ *K, and the undecided area may converge to a nonzero limit supported on {B* ≤ 0 *< φ}*.

*Refining* [**p**] *at ε* = 0.05 *changes the certified area by less than* 0.02 *percentage points at approximately* 100*× the runtime. We report the fixed-*[**p**] *variant in the numerical experiments and use the refining form only to establish completeness in principle. The non-degeneracy hypothesis is assumed, not verified, for* (1); *the empirical relation* |*U* (*ε*)| *≈* 0.35 *ε is consistent with it over the range explored*.

#### Remark 3

(Factored versus expanded evaluation). *The factored Routh expression a*_1_(*a*_2_*a*_3_−*a*_1_*a*_4_)− *a encloses D*_3_ *with substantially fewer repeated variable occurrences than the expanded polynomial q. Under the same outward-rounded natural interval extension and on the same paving, the factored evaluation certifies a larger area than the expanded evaluation at roughly half the runtime (Table 1)*. *Since the two enclosures apply to different functions, namely D*_3_ *and* 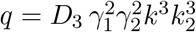, *their interval widths are not directly comparable; the comparison of interest is between the resulting certified regions*.

*The expansion to q is introduced not to improve tightness, but to obtain the coefficient representation required by the Bernstein enclosure. In settings where clearing denominators substantially increases degree or term count (e*.*g*., *Hill-type kinetics), a factored evaluation may remain preferable; such cases are not pursued here*.

**Table 1:** Certified area, search size, and runtime for five tests on an identical paving, design domain, witness set, and stopping rule. **Boxes** counts the number of terminal boxes in the final subpaving. Rows 1–4 correspond to: (1) the natural interval extension of *q*; (2) a centered (mean-value) form of *q*; (3) the natural extension applied to the unexpanded Routh expression (Remark 3); and (4) a Kharitonov test with coefficient intervals computed over a 64^3^ subdivision of [**p**], with exclusion certified by the same witness test as row 1. Times are obtained in outward-rounded arithmetic for rows 1–3 and in floating point for the Bernstein rows; the outward-rounded verification cost for the Bernstein rows is reported in Section 5.8.

| Test | $\varepsilon$ | Boxes | Certified | Excluded | Undecided | Time |
| --- | --- | --- | --- | --- | --- | --- |
| natural extension | 0.05 | 3,897 | 50.93% | 10.80% | 38.27% | 73.9 s |
| centered form | 0.05 | 3,729 | 50.93% | 14.01% | 35.06% | 546.4 s |
| factored form | 0.05 | 3,455 | 52.86% | 11.72% | 35.42% | 36.0 s |
| Kharitonov | 0.05 | 3,017 | 58.86% | 10.80% | 30.33% | 408.6 s |
| Bernstein | 0.10 | 158 | 82.08% | 14.40% | 3.52% | 0.3 s |
| Bernstein | 0.05 | 324 | 83.19% | 15.03% | 1.78% | 0.7 s |
| Bernstein | 0.025 | 666 | 83.71% | 15.37% | 0.92% | 1.4 s |

## 4 Algorithm

All arithmetic is outward-rounded, so each classified terminal box is accompanied by a machine-checkable certificate. Design-box endpoints are outward-rounded to enclose the intended domain. When the plant parameters are specialized to a corner, the Bernstein evaluation reduces to a lower-dimensional tensor obtained by specializing those coordinates, so the witness check can be implemented by contracting the coefficient tensor along the plant axes.

## 5 Numerical Results

Sections 5.1–5.6 treat the running example (1), and Section 5.7 a fifth-order loop. We take *γ*_1_ ∈ [0.8, 1.2], *γ*_2_ ∈ [0.4, 0.6] and *k*_2_ ∈ [0.9, 1.1], uncertainties of *±*20%, *±*20% and *±*10% respectively, with *µ* = *θ* = 1. Concentrations are reported in nM and first-order rates in h^−1^ ; the sequestration rate *η* has units nM^−1^h^−1^ and *µ* has units nM h^−1^. The uncertainty levels (*±*10– 20%) are comparable to the accuracy with which such kinetic constants are typically reported [17]. We pave the design plane over log_10_ *η* ∈ [−2, 2] and log_10_ *k* ∈ [−1, 2]; all areas reported below are fractions of this log-rectangle *D*. Unless stated otherwise, baseline pavings are computed entirely in outward-rounded arithmetic. Bernstein pavings are computed in floating point and verified by outward-rounded recomputation (Section 5.8); the verified classifications constitute the certificates reported below. Timings are single-core on a 2.8 GHz Intel Xeon.

### Algorithm 1

Certified design region

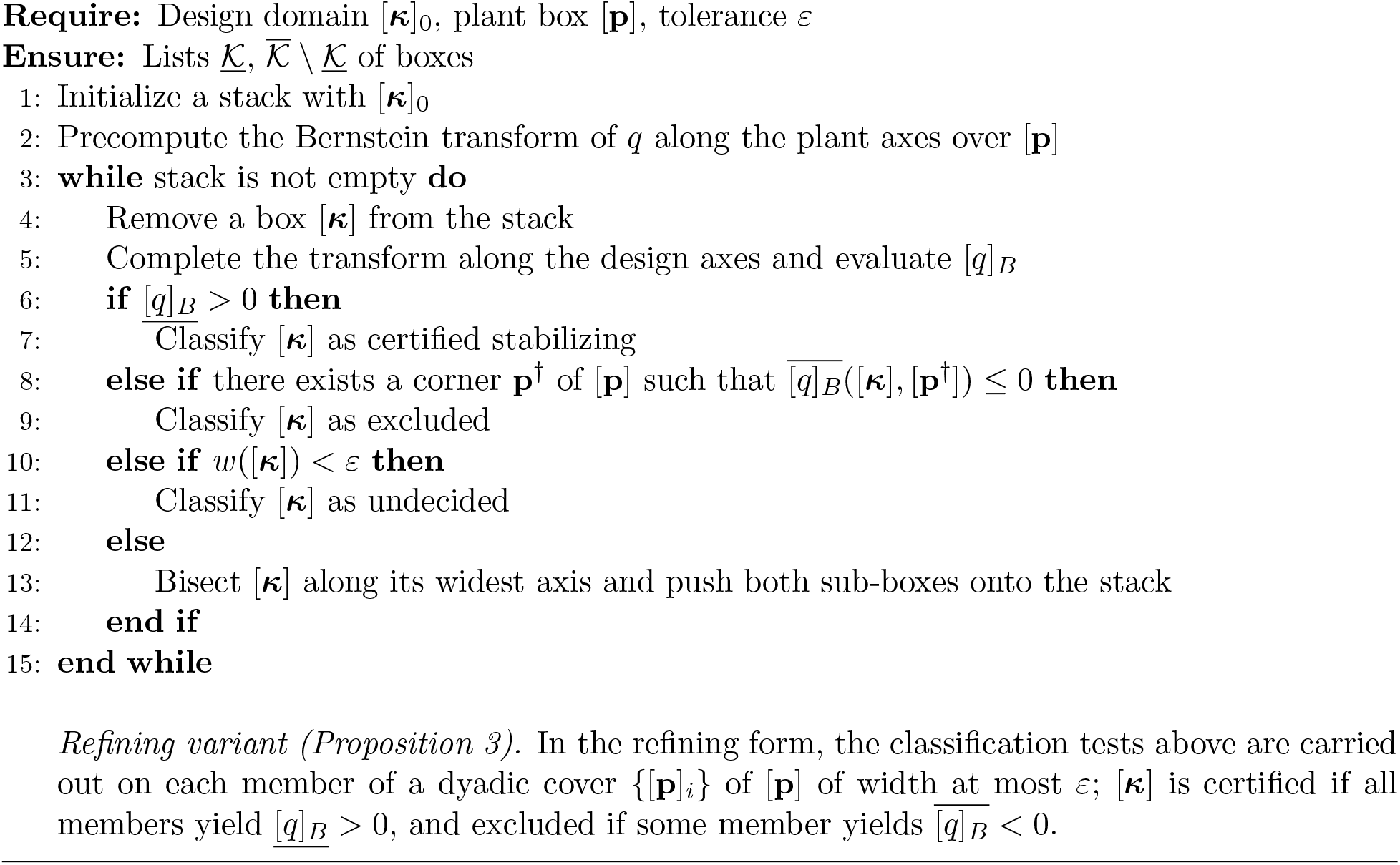

### 5.1 Certified and sampled design regions

To quantify the conservatism of the certified regions, we compute a sampling-based reference estimate on a dense design grid. Specifically, on a 400 *×* 400 grid in *D* we evaluate *D*_3_ at the 2^3^ corners of [**p**] and at 200 additional interior samples per grid point; repeating the computation on an 800 *×* 800 grid changes the resulting area estimate by at most 0.01 percentage points. For a finite plant sample set *P*_*s*_, the sampled acceptance set *K*_*s*_ = *{****κ*** : *q*(***κ*, p**) *>* 0 ∀ **p** ∈ *P*_*s*_*}* satisfies *K* ⊆ *K*_*s*_, i.e., plant sampling can only miss violations. The sampled area is therefore reported only as a reference estimate of |*K*_*s*_|*/*|*D*|. Since *D*_3_ is not monotone in **p**, corner evaluation alone would not yield a sound robust test. A rigorous two-sided bracket is instead obtained directly from the certified paving (Section 5.3).

Table 1 and Fig. 1 summarize the result. With the natural interval extension, 50.93% of the design plane is certified, whereas the sampled reference estimate is 84.13%. The undecided boxes in Fig. 1(a) correspond to regions in which the range enclosure is too conservative to determine the sign of *q*; the indeterminacy arises from the enclosure rather than from the underlying dynamics. The natural extension is the least tight among the standard interval evaluations, so we also consider two strengthened enclosures on the identical paving. The centered (mean-value) form *q*(**x**) ∈ *q*(**c**) + ∑_*i*_[*∂q/∂x*_*i*_]([**x**]) ([**x**]_*i*_ −**c**_*i*_) about the midpoint **c** of [**x**], intersected with the natural extension, has excess width *O*(*w*^2^) rather than *O*(*w*). At *ε* = 0.05 (Table 1), it certifies the same stabilizing area as the natural extension, while increasing the excluded fraction, at approximately seven times the runtime. The natural extension applied to the unexpanded Routh expression (Remark 3) yields a modest improvement, certifying 52.86% at *ε* = 0.05 and 54.12% at *ε* = 0.025. Both remain substantially below the Bernstein paving.

**Figure 1:**
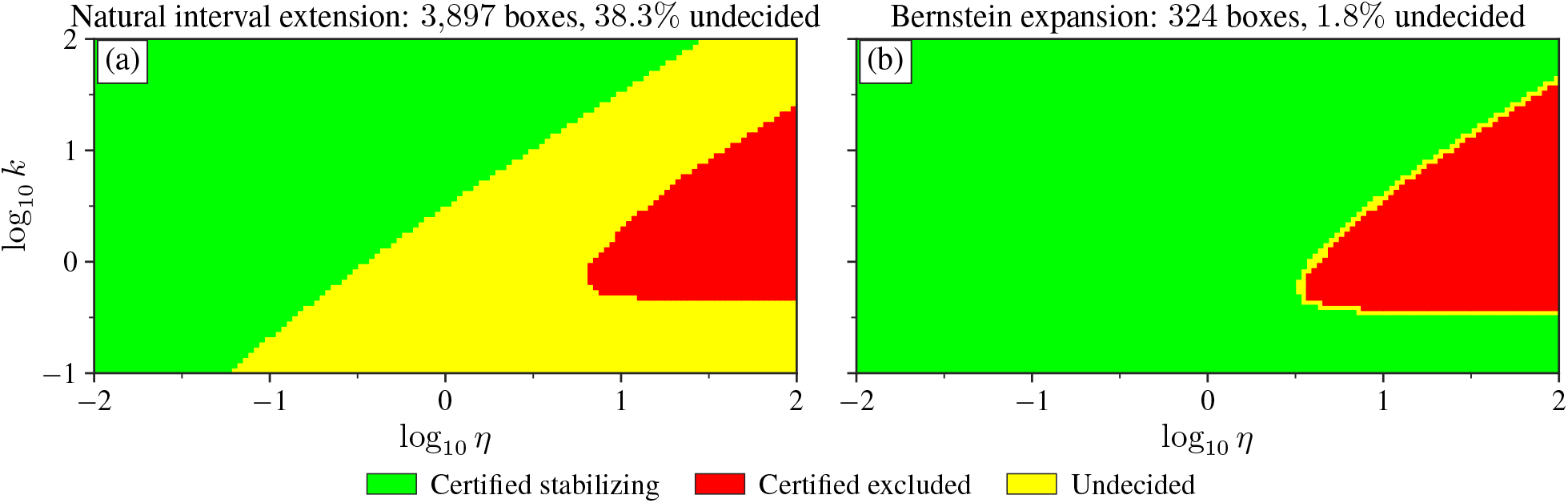
Certified subpavings of the design plane for *γ*_1_ ∈ [0.8, 1.2], *γ*_2_ ∈ [0.4, 0.6], *k*_2_ ∈ [0.9, 1.1], *µ* = *θ* = 1, both panels at the same bisection tolerance *ε* = 0.05. (a) Natural interval extension of *q*; (b) Bernstein range bounds, where the undecided set is confined to a thin band near the stability boundary. Exclusion is certified in both panels using a witness plant.

Kharitonov’s theorem gives a fourth baseline that does not bound *q* at all: enclosing *a*_1_, …, *a*_4_ of (4) in intervals over [***κ***] *×* [**p**] and requiring the four Kharitonov polynomials to be Hurwitz is a sound inner test. With the coefficient intervals evaluated directly in interval arithmetic, the test certifies only 43.44% at *ε* = 0.05. To separate the conservatism of the theorem from that of the coefficient enclosure, we recomputed the coefficient intervals over a 64^3^ subdivision of [**p**], where going from 32^3^ to 64^3^ changes the certified area by only 0.35 percentage points. The certified area then rises to 58.86% (Table 1), still about 24 percentage points below the Bernstein paving. The remaining gap comes from the theorem itself, which treats *a*_1_, …, *a*_4_ as independent although they share parameters. As a result, the coefficient box defines a larger interval-polynomial family than the one induced by the coupled parameters (***κ*, p**), and the test can be conservative even at single design points (accepting only about 61% of a 200 *×* 200 grid).

The gap is not an artifact of leaving the plant box unrefined or of the choice of *D*. Subdividing [**p**] into 2^3^, 3^3^, and 4^3^ parts and taking the minimum of the lower bounds raises the natural extension to 59.62%, 63.16%, and 65.26%, at about six times the runtime for 4^3^, still some 18 percentage points below the Bernstein paving. Repeating the *ε* = 0.05 comparison on other design domains gives 52.45% versus 84.41% on log_10_ *η* ∈ [−3, 3], log_10_ *k* ∈ [−2, 3] and 74.49% versus 96.41% on log_10_ *η* ∈ [−1, 1], log_10_ *k* ∈ [0, 2]; the absolute gap depends on *D*, but the Bernstein paving stays close to the sampled reference estimate in both cases.

Replacing the bound by (7) certifies 83.71% at *ε* = 0.025, with an undecided area of 0.92%. By the two-sided paving bracket of Section 5.3, this implies the (unknown) robustly stabilizing area lies in [83.70%, 84.63%] on the stated design domain. At a common tolerance *ε* = 0.05, the certified area increases from 50.93% to 83.19% and the number of terminal boxes decreases from 3,897 to 324. The runtimes are not directly comparable: the natural-extension paving is computed entirely in outward-rounded arithmetic (73.9 s), whereas the Bernstein paving is computed in floating point (0.7 s) and then verified in outward-rounded arithmetic (111.1 s; Section 5.8). The total time is therefore larger in the Bernstein pipeline for this example. Fig. 2 reports the dependence on *ε*. The closed form (6) offers an independent check of the certified paving. *In contrast to this asymptotic calculation, the certified paving localizes the stability boundary over the full design domain and yields a rigorous two-sided bracket on the stabilizing area at the chosen tolerance*. Among the terminal boxes in Fig. 1(b) that touch log_10_ *η* = 2, the certified ones end at log_10_ *k* = −0.484375 and the excluded ones begin at −0.4375, which bracket the robust threshold −0.457 derived in Section 2.2.

**Figure 2:**
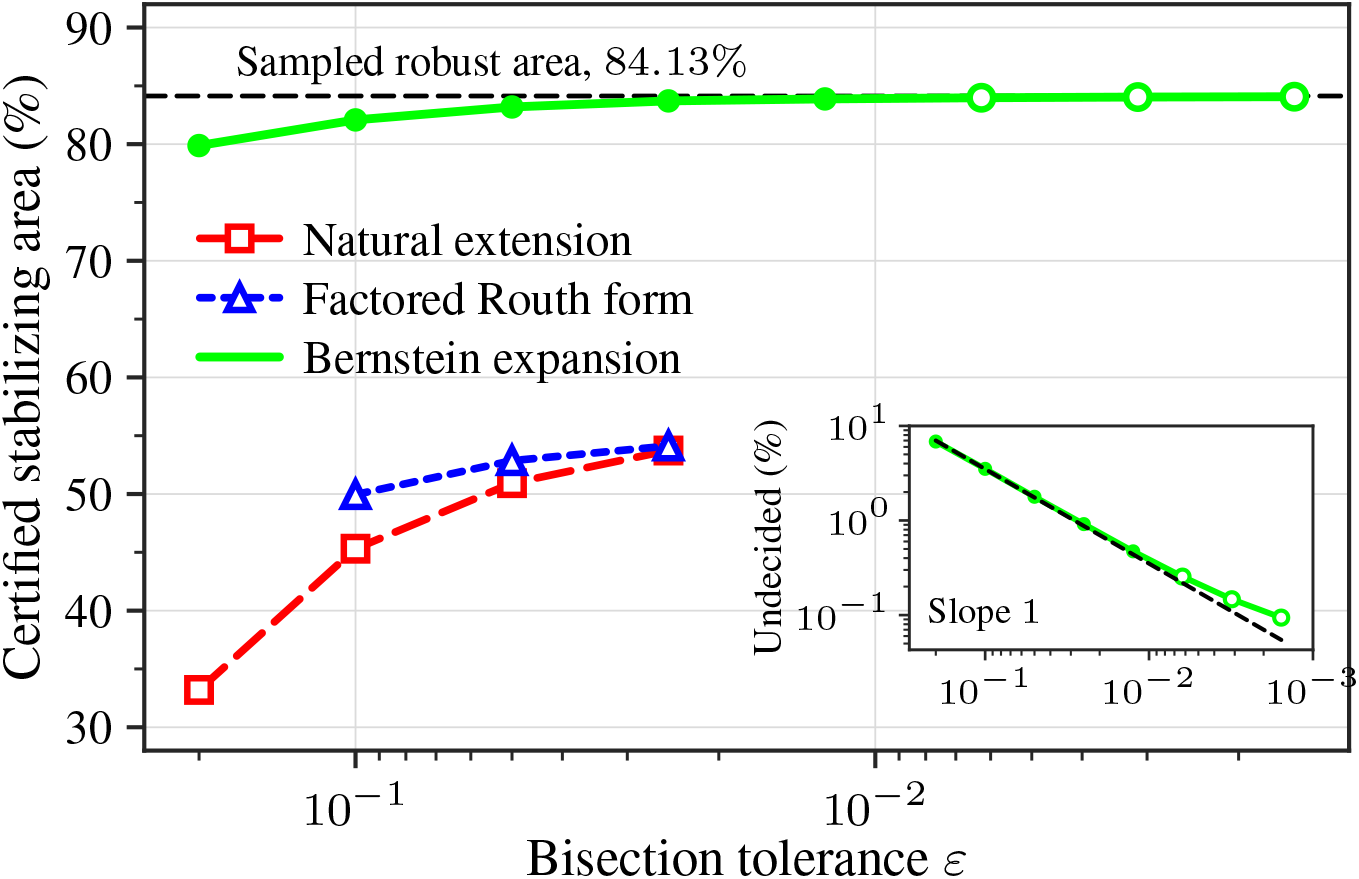
Certified stabilizing area versus the bisection tolerance *ε* for the natural interval extension, the factored Routh form, and the Bernstein enclosure (the centered form coincides with the natural extension and is omitted). Filled Bernstein markers correspond to outward-rounded computations; hollow markers (*ε* ≤ 6.25 *×* 10^−3^) correspond to unverified floating-point runs. The baseline curves are truncated to limit runtime. Inset: undecided area of the Bernstein paving, approximately linear in *ε*. Since [**p**] is held fixed, the inset reports an empirical trend and does not constitute a test of Proposition 3.

Applying the 30-point spot check yields no disagreements with the classifications.

### 5.2 Cost of nominal tuning

Evaluating *D*_3_ at the nominal plant (*γ*_1_, *γ*_2_, *k*_2_) = (1, 0.5, 1) labels 89.62% of the design domain as stabilizing, whereas the sampling-based reference estimate of the robustly stabilizing fraction is 84.13% (Section 5.1). Since the nominal plant is not included in the sample set *P*_*s*_, the corresponding accepted sets need not be nested. On the reference grid, 5.49% of design points are accepted at the nominal plant and rejected by *P*_*s*_, and none are rejected at the nominal plant but accepted by *P*_*s*_. The certified paving provides a rigorous counterpart to this discrepancy. Summing terminal boxes that (i) satisfy the nominal condition [*q*]_*B*_([***κ***], [**p**_nom_]) *>* 0 and (ii) are excluded by Proposition 1(2), at least 2.60% of the design domain at *ε* = 0.05 (and 2.94% at *ε* = 0.025) is certified stabilizing at the nominal plant but certified not robustly stabilizing. These lower bounds increase as the paving is refined.

This effect is illustrated by a representative terminal box. At *ε* = 0.05, the box log_10_ *η* ∈ [1.75, 2], log_10_ *k* ∈ [−0.4375, −0.25] is certified stabilizing at the nominal plant, with Bernstein lower bound 1.1 *×* 10^3^, but every gain in the box is destabilizing for the admissible corner (*γ*_1_, *γ*_2_, *k*_2_) = (0.8, 0.4, 1.1). The box lies in the band −0.457 *<* log_10_ *k <* −0.125 between the robust and nominal thresholds of Section 2.2. By contrast, the single box log_10_ *η* ∈ [−2, 0], log_10_ *k* ∈ [−1, 2] is certified stabilizing for the full uncertainty set and is returned as a single certificate rather than as a sampled region.

### 5.3 Convergence

Table 2 reports the Bernstein paving under successive halvings of *ε*. The undecided fraction decreases by approximately a factor of two at each step, and the relation |*U* (*ε*)| *≈* 0.35 *ε* provides a convenient empirical summary over the range shown. At the smallest tolerances the ratio |*U* (*ε*)|*/*|*U* (*ε/*2)| drops below two, consistent with the saturation mechanism described in Remark 2. A fit over finitely many tolerances does not, however, establish a nonzero asymptotic lower bound on |*U* (*ε*)|.

**Table 2:** Convergence of the Bernstein test. Rows down to *ε* = 1.25 *×* 10^−2^ were recomputed with outward-rounded arithmetic (Section 5.8); for smaller *ε*, values are unverified floating-point results. The last column reports |*U* (*ε*)|*/*|*U* (*ε/*2)| between consecutive rows. **Boxes** counts the number of terminal boxes in the final subpaving.

| $\varepsilon$ | Boxes | Certified | Excluded | Undecided | Ratio |
| --- | --- | --- | --- | --- | --- |
| $2.0 \times 10^{-1}$ | 76 | 79.88% | 13.28% | 6.836% | — |
| $1.0 \times 10^{-1}$ | 158 | 82.08% | 14.40% | 3.516% | 1.94 |
| $5.0 \times 10^{-2}$ | 324 | 83.19% | 15.03% | 1.782% | 1.97 |
| $2.5 \times 10^{-2}$ | 666 | 83.71% | 15.37% | 0.916% | 1.95 |
| $1.25 \times 10^{-2}$ | 1369 | 83.88% | 15.65% | 0.472% | 1.94 |
| $6.25 \times 10^{-3}$ | 2839 | 83.98% | 15.77% | 0.255% | 1.85 |
| $3.13 \times 10^{-3}$ | 6084 | 84.04% | 15.81% | 0.147% | 1.73 |
| $1.56 \times 10^{-3}$ | 13820 | 84.07% | 15.84% | 0.094% | 1.57 |

The certified paving also yields a rigorous two-sided bracket on the robustly stabilizing area. Let *C* and *E* denote the certified-stabilizing and certified-excluded regions, respectively. By soundness, *C* ⊆ *K* ⊆ *D \ E*, and therefore |*C*| ≤ |*K*| ≤ |*D*| − |*E*|. At *ε* = 0.025 this gives 83.70% ≤ |*K*|*/*|*D*| ≤ 84.63%, i.e., a window of 0.93 percentage points obtained without sampling; this row was recomputed and verified in outward-rounded arithmetic (Section 5.8).

Since terminal boxes are dyadic in the bisected coordinates, their areas (and hence the totals |*C*| and |*E*|) are accumulated exactly; rounding occurs only in the displayed percentages. The two bracket endpoints are rounded outward from 83.7097% and 84.6252%, whereas the entries of Tables 1 and 2 are rounded to nearest. The sampled value 84.13% lies within the certified bracket and is retained only as a reference estimate.

### 5.4 Tolerable uncertainty levels

We apply a uniform relative uncertainty *±w* to all three plant parameters at *ε* = 0.05. The case *w* = 20% therefore differs from Table 1, which uses *±*10% on *k*_2_. As *w* increases from 5% to 40%, the certified stabilizing area decreases from 87.28% to 72.61% and the number of terminal boxes increases from 279 to 434. Over the same range, the gap between certification and the sampling-based reference estimate increases from 0.85 to 1.47 percentage points. By contrast, nominal evaluation remains at 89.62% for all *w*, and the nominal–sampled discrepancy on the reference grid increases from 1.49 to 15.54 percentage points.

### 5.5 Sampling limitations

We assess whether dense Monte Carlo sampling can serve as a practical surrogate for certification. On a 400 *×* 400 design grid (160,000 points), we draw *N* plants independently and uniformly from [**p**] (fixed seed <monospace>MersenneTwister(5)</monospace> in the released code; corners excluded) and accept a gain if none of the *N* sampled plants destabilizes it. Using the reference set *K*_*s*_ from Section 5.1 (which does include the corners), the fraction of design points accepted by this rule but rejected by the reference is 6.82% at *N* = 1, 2.41% at *N* = 10, 1.44% at *N* = 100, and 0.55% at *N* = 1000; the latter requires 1.6 *×* 10^8^ determinant evaluations. Since *K*_*s*_ is itself based on a finite plant sample, these values are lower bounds on the true false-accept rate. The error decreases with *N*, but sampling does not yield a certificate of robust stability.

### 5.6 A third gain

Set inversion scales exponentially with the number of design parameters. Allowing the sensing gain *θ* to vary makes the design space three dimensional, and clearing denominators yields 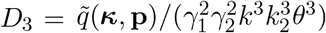, where the denominator is a strictly positive monomial. The polynomial 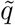 has multidegree (7, 7, 6) in (*γ*_1_, *γ*_2_, *k*_2_) and (3, 6, 6) in (*η, k, θ*), corresponding to 87,808 Bernstein tensor entries (versus 12,544 in the two-gain case).

Paving log_10_ *θ* ∈ [−1, 1] in addition yields certified stabilizing volume fractions of 80.52% at *ε* = 0.2, 82.73% at *ε* = 0.1, and 83.77% at *ε* = 0.05, using 1,191, 4,814, and 19,532 terminal boxes. These results are based on floating-point classifications and are reported as a scaling study; the reported volume fractions should therefore be interpreted as empirical.

### 5.7 A higher-order plant

We append 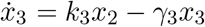 to (1) and let the controller sense *x*_3_. The closed loop is fifth order, and its five characteristic coefficients are again sums of positive monomials; by the Liénard–Chipart criterion, stability is equivalent to Δ_2_ *>* 0 and Δ_4_ *>* 0 for the Hurwitz determinants Δ_*j*_ . After substitution, these conditions are polynomials in (*γ*_1_, *γ*_2_, *γ*_3_, *η, k*) of multidegree (5, 5, 5, 2, 4) and (10, 10, 10, 4, 8) with 28 and 154 terms; the larger tensor has 59,895 entries. All coefficients of Δ_2_ are positive, so Δ_2_ *>* 0 holds throughout the positive orthant; nevertheless, we retain both conditions Δ_2_ *>* 0 and Δ_4_ *>* 0 so that the example instantiates Definition 1 with *m* = 2.

The asymptotic argument of Section 2.2 also extends to this case. The leading coefficient of Δ_4_ in *η* is a positive multiple of *γ*_1_*γ*_2_*γ*_3_(*γ*_1_ + *γ*_2_)(*γ*_2_ + *γ*_3_)(*γ*_1_ + *γ*_3_) − (*γ*_1_ + *γ*_2_ + *γ*_3_)^2^*k*. The resulting threshold on *k* increases with each *γ*_*i*_, so over the uncertainty box the robust threshold log_10_ *k <* −0.439 is attained at the corner (0.8, 0.8, 0.8); the corresponding nominal threshold is log_10_ *k <* −0.051.

We take *±*20% uncertainty on the three degradation rates, fix *k*_2_ = *k*_3_ = 1, and pave the same design plane at *ε* = 0.05. A design box is excluded when either Δ_2_ or Δ_4_ is non-positive over it at one of the 2^3^ corners of [**p**], as in Proposition 1(2). The natural interval extension certifies 34.44% of the plane and excludes 18.35%, leaving 47.22% undecided with 4,763 terminal boxes. The Bernstein bound certifies 73.79% and excludes 24.10%, leaving 2.11% undecided with 389 terminal boxes; consequently, the robustly stabilizing area lies in [73.79%, 75.91%]. The sampling-based reference estimate 74.87% falls within this bracket. All 216 classified (certified or excluded) Bernstein terminal boxes retain their floating-point classification under outward-rounded recomputation. Among the terminal boxes touching log_10_ *η* = 2, the certified ones extend up to log_10_ *k* = −0.484375 and the excluded ones begin at −0.4375, bracketing the robust threshold −0.439. The same 30-point spot check yields no disagreements with the classifications.

For timing comparisons, exclusion was disabled (the certified areas are unchanged). The natural-extension paving required 117.0 s in outward-rounded arithmetic. The Bernstein computation required 3.8 s in floating point plus 247.9 s for outward-rounded verification, for a total of 251.7 s. Nominal evaluation at (*γ*_1_, *γ*_2_, *γ*_3_) = (1, 1, 1) marks 82.80% of the plane as stabilizing, a 7.93-percentage-point difference relative to the sampling-based reference estimate. Although two examples do not establish a systematic dependence on system order, they suggest that the improvement obtained with Bernstein range bounds is not specific to (1).

### 5.8 Verification in exact arithmetic

The Bernstein transform involves binomial coefficients and powers of interval endpoints spanning several decades. The associated basis changes can therefore suffer from cancellation in floating-point arithmetic. We therefore recomputed the core pavings using outward-rounded arithmetic throughout (binomial coefficients; affine maps and basis-change matrices; coefficient accumulations; and design-box endpoints 10^*e*^, widened outward to account for non-representability). Every Bernstein terminal box retained its floating-point classification under this recomputation. Consequently, the Bernstein rows of Table 1, the uncertainty sweep of Section 5.4, the fifth-order runs of Section 5.7, and the entries of Table 2 down to *ε* = 1.25 *×* 10^−2^ are reproduced exactly. The three finest rows of Table 2 and the three-gain results of Section 5.6 were not recomputed with outward-rounded arithmetic and are reported as unverified floating-point results. The outward-rounded recomputation is substantially more expensive: for the *ε* = 0.025 Bernstein paving it costs 232 s versus 1.4 s in floating point, motivating a separate verification phase. Moreover, we re-derived in exact rational arithmetic the twelve tightest certified-stabilizing boxes (verifying min_*α*_ *b*_*α*_ *>* 0 over [**p**]) and the eight tightest excluded boxes (verifying that some plant corner yields max_*α*_ *b*_*α*_ ≤ 0). All twenty checks succeed, with the smallest margins 9.1 *×* 10^−4^ and 3.0 *×* 10^−4^.

#### Reproducibility

Since the reported regions are certificates, they should be independently checkable. The code used to generate all tables and figures is available online [18].

## 6 Conclusions

We formulated controller tuning under parametric plant uncertainty as a universally quantified set-inversion problem and computed inner and outer box approximations of the gain set that stabilizes every plant in a prescribed uncertainty box. The resulting certificates guarantee local exponential stability of the deterministic equilibrium for each fixed admissible parameter vector; they do not address time-varying uncertainty or stochastic dynamics. In the two examples considered, robust stability reduces to polynomial sign conditions that can be evaluated with validated range enclosures. The numerical results indicate that the tightness of the range enclosure has a substantial effect on the certified stabilizing region: for both the fourth- and fifth-order loops, Bernstein range bounds certify larger stabilizing regions than standard interval extensions and a Kharitonov baseline. The remaining undecided set yields a rigorous two-sided bracket on the stabilizing area at the chosen tolerance. The computational cost increases with the number of free controller parameters, motivating structure-exploiting implementations and improved search strategies. The present formulation also relies on an explicitly available equilibrium. For models with implicit equilibria, such as Hill-type kinetics, one may combine equilibrium enclosure with stability certification while enforcing the equilibrium manifold as a constraint rather than enclosing it by an unconstrained box.

## References

[1] C. Briat, A. Gupta, and M. Khammash, “Antithetic integral feedback ensures robust perfect adaptation in noisy biomolecular networks,” Cell systems, vol. 2, no. 1, pp. 15–26, 2016.

[2] S. K. Aoki, G. Lillacci, A. Gupta, A. Baumschlager, D. Schweingruber, and M. Khammash, “A universal biomolecular integral feedback controller for robust perfect adaptation,” Nature, vol. 570, no. 7762, pp. 533–537, 2019.

[3] N. Olsman, A.-A. Baetica, F. Xiao, Y. P. Leong, R. M. Murray, and J. C. Doyle, “Hard limits and performance tradeoffs in a class of antithetic integral feedback networks,” Cell systems, vol. 9, no. 1, pp. 49–63, 2019.

[4] L. Jaulin, M. Kieffer, O. Didrit, and E. Walter, “Interval analysis,” in Applied Interval Analysis: With Examples in Parameter and State Estimation, Robust Control and Robotics. Springer, 2001, pp. 11–43.

[5] R. E. Moore, R. B. Kearfott, and M. J. Cloud, Introduction to interval analysis. SIAM, 2009.

[6] R. Prakash, S. Janardhanan, and S. Sen, “Comparing point and interval methods for equilibrium computation under parametric uncertainty,” arXiv preprint 2608.07071, 2026.

[7] R. Prakash, S. Janardhanan, and S. Sen, “Certified detection of bifurcation candidates in uncertain nonlinear systems using interval analysis,” arXiv preprint 2608.07119, 2026.

[8] R. Prakash, S. Janardhanan, and S. Sen, “Computational complexity analysis of interval methods in solving uncertain nonlinear systems,” arXiv preprint 2603.19965, 2026.

[9] J. Ackermann, “Parameter space design of robust control systems,” IEEE Transactions on Automatic Control, vol. 25, no. 6, pp. 1058–1072, 1980.

[10] S. P. Bhattacharyya, H. Chapellat, and L. H. Keel, Robust Control: The Parametric Approach. Prentice Hall PTR, 1995.

[11] M. Zettler and J. Garloff, “Robustness analysis of polynomials with polynomial parameter dependency using bernstein expansion,” IEEE Transactions on Automatic Control, vol. 43, no. 3, pp. 425–431, 1998.

[12] J. Garloff, “Application of bernstein expansion to the solution of control problems,” Reliable computing, vol. 6, no. 3, pp. 303–320, 2000.

[13] S. Malan, M. Milanese, and M. Taragna, “Robust analysis and design of control systems using interval arithmetic,” Automatica, vol. 33, no. 7, pp. 1363–1372, 1997.

[14] P. A. Parrilo, “Semidefinite programming relaxations for semialgebraic problems,” Mathematical programming, vol. 96, no. 2, pp. 293–320, 2003.

[15] É. Walter and L. Jaulin, “Guaranteed characterization of stability domains via set inversion,” IEEE Transactions on Automatic Control, vol. 39, no. 4, pp. 886–889, 1994.

[16] R. Prakash, S. Janardhanan, and S. Sen, “Design of parameter intervals to meet steady-state specifications in biomolecular circuits using interval analysis,” bioRxiv, doi: 10.1101/2024.10.24.620126, 2024.

[17] R. Milo, P. Jorgensen, U. Moran, G. Weber, and M. Springer, “Bionumbers—the database of key numbers in molecular and cell biology,” Nucleic acids research, vol. 38, no. Suppl 1, pp. D750–D753, 2010.

[18] R. Prakash, “Code for examples,” https://github.com/RudraPrakashIITDelhi/AntitheticIntegralFeedback, 2026, accessed: 13 September 2026.

